# A Cas9 with Complete PAM Recognition for Adenine Dinucleotides

**DOI:** 10.1101/429654

**Authors:** Noah Jakimo, Pranam Chatterjee, Lisa Nip, Joseph M Jacobson

## Abstract

CRISPR-associated (Cas) DNA-endonucleases are remarkably effective tools for genome engineering, but have limited target ranges due to their protospacer adjacent motif (PAM) requirements. We demonstrate a critical expansion of the targetable sequence space for a Type-IIA CRISPR-associated enzyme through identification of the natural 5’-NAA-3’ PAM specificity of a *Streptococcus macacae* Cas9 (Smac Cas9). We further recombine protein domains between Smac Cas9 and its well-established ortholog from *Streptococcus pyogenes* (Spy Cas9), as well as an “increased” nucleolytic variant (iSpy Cas9), to achieve consistent mediation of gene modification and base editing. In a comparison to previously reported Cas9 and Cas12a enzymes, we show that our hybrids recognize all adenine dinucleotide PAM sequences and possess robust editing efficiency in human cells.

---

Biotechnologies based on RNA-guided CRISPR systems have enabled precise and programmable genomic interfacing.^1^ However, CRISPR-associated (Cas) endonucleases are also collectively restrained from localizing to any position along double-stranded DNA (dsDNA) due to their requirement for targets to neighbor a protospacer adajacent motif (PAM).^2–4^ Current gaps in the PAM sequences that Cas enzymes are known to recognize prevent access to numerous genomic positions for powerful methods like base editing, which can only operate on a narrow window of nucleotides at fixed distances from the PAM.^5^ Many AT-rich regions, in particular, have been excluded from compelling CRISPR applications because previously reported endonucleases, such as Cas9 and Cas12a (formerly known as Cpf1), require targets to neighbor GC-content or more restrictive motifs, respectively.^6–8^

In this work, we introduce a Cas9 ortholog derived from *Streptococcus macacae NCTC 11558* that can instead recognize a short 5’-NAA-3’ PAM.^9^ These sequences constitute 18.6% of the human genome, making adjacent adenines the most abundant dinucleotide (Supplementary Figure S1A-B). The importance of this alternative PAM recognition for a Cas9 enzyme is reinforced by recent work exposing that many Cas12a orthologs, while targeting dsDNA at AT-rich PAM sites with intrinsic high fidelity, will indiscriminately digest single-stranded DNA (ssDNA) once bound to their targets.^10,11^ Such collateral activity may introduce unwanted risks around partially unpaired chromosomal structures, such as transcription bubbles, R-loops, and replication forks. Here we present engineered nucleases derived from Smac Cas9 and characterize their novel specificity and utility by means of transcriptional repression in bacterial culture, *in vitro* digestion reactions, and both gene and base editing in a human cell line.

To modify the ancestral 5’-NGG-3’ PAM specificity of Spy Cas9, early and new reports have employed directed evolution (e.g., “VQR”, “EQR”, and “VRER” variants) and rational design informed by crystal structure (e.g., “QQR” and “NG” variants).^12–15^ These reports focused on the PAM-contacting arginine residues R1333 and R1335 that abolish function when exclusively mutated. While those studies identified compensatory mutations resulting in altered PAM specificity, the Cas9 variants that they produced maintained a guanine preference in at least one position of the PAM sequence for reported *in vivo* editing. We aimed to lift such GC-content pre-requisites via a custom bioinformatics-driven workflow that mines existing PAM diversity in the *Streptococcus* genus. Using that workflow we homed in on Smac Cas9 as having the potential to bear novel PAM specificity upon aligning 115 orthologs of Spy Cas9 from UniProt (limited to those with greater than a 70% pairwise BLOSSOM62 score). From the alignment we found Smac Cas9 was one of two close homologs, along with a *Streptococcus mutans B112SM-A* Cas9 (Smut Cas9), with divergence at both of the positions aligned to the otherwise highly conserved PAM-contacting arginines (Figure 1A-B; Supplementary Figure S2A). We thus hypothesized that Smac Cas9 had naturally co-evolved the necessary compensatory mutations to gain new PAM recognition. A small sample size of 13 spacers from its corresponding genome’s CRISPR cassette prevented us from confidently inferring the Smac Cas9 PAM *in silico*. However, the possibility for Smac Cas9 requiring less GC-content in its PAM was supported by sequence similarities to the “QQR” variant that has 5’-NAAG-3’ specificity, in addition to the AT-rich putative consensus PAM for phage-originating spacers in CRISPR cassettes associated with highly homologous Smut Cas9, which were identified with the aid of our computational pipeline called SPAMALOT (Figure 1C; Supplementary Figure S2B; Supplementary Figure S3).^16^

**Figure 1:**
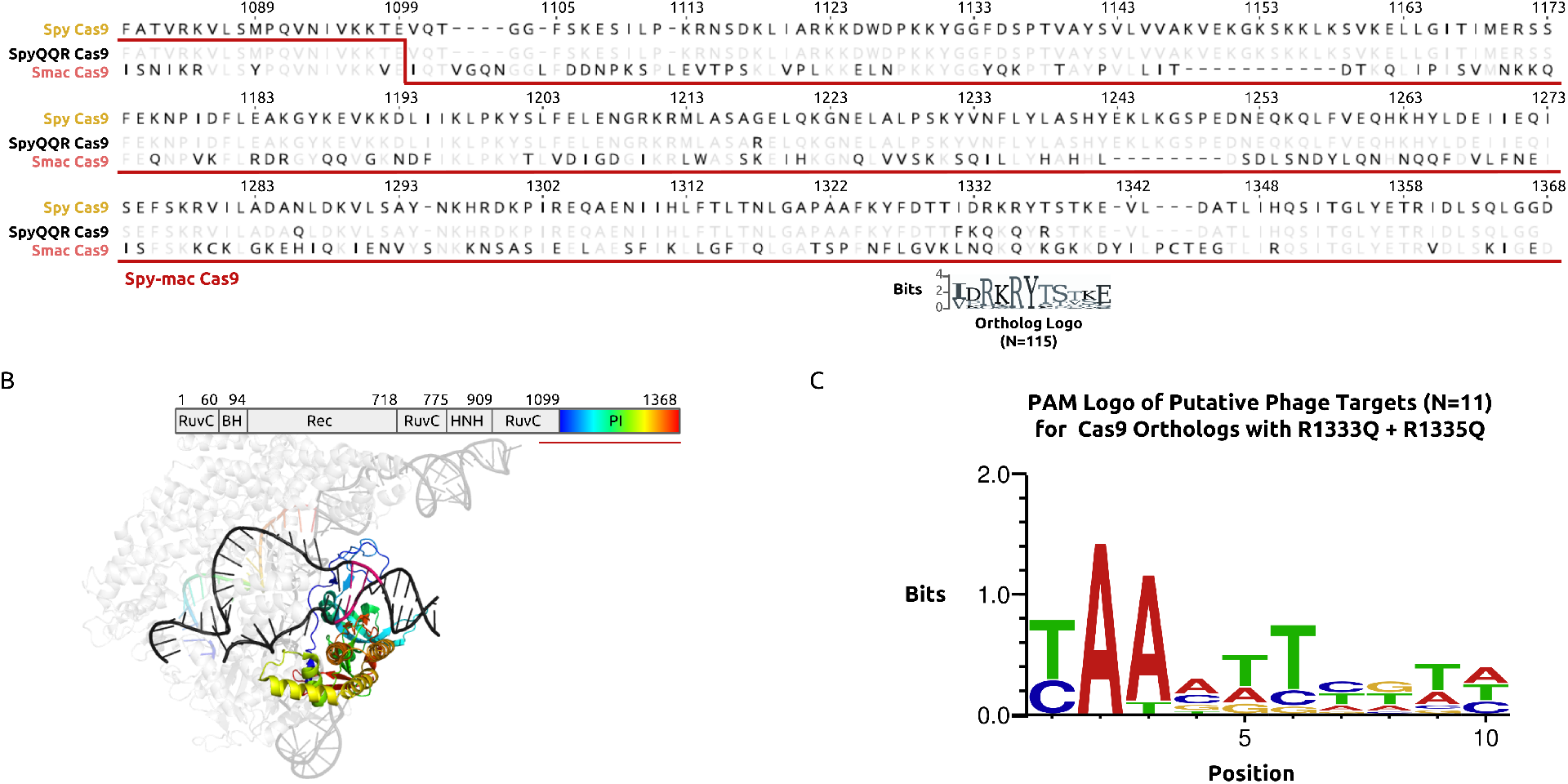
Identification of features from natural PAM divergence through bioinformatics. **(A)** Sequence alignment (Genewiz software) of Spy Cas9, its “QQR” variant, and Smac Cas9. The step in the underlining red line marks the joining of Spy Cas9 and Smac Cas9 to construct a Spy-mac Cas9 hybrid. The sequence logo (Weblogo online tool) immediately below the alignment depicts the conservation at 11 positions around the PAM-contacting arginines of Spy Cas9. **(B)** The domain organization of Spy Cas9 juxtaposed over a color-coded structure of RNA-guided, target-bound Spy Cas9 (PDB ID 5F9R). The two DNA strands are black with the exception of a magenta segment corresponding to the PAM. A blue-green-red color map is used for labeling the Cas9 PI domain and guide spacer sequence to highlight structures that confer sequence specificity and the prevalence of intra-domain contacts within the PI.^42^ **(C)** A sequence logo generated online (WebLogo) that was input with putative PAM sequences found in *Streptococcus phage* and associated with close Smac Cas9 homologs.

We proceeded to experimentally assay the PAM preferences of several *Streptococcus* orthologs that change one or both of the critical PAM-contacts. Based on demonstrated examples of the PAM-interaction (PI) domain and guide RNA (gRNA) having cross-compatibility between Cas9 orthologs that are closely related and active, we constructed new variants by rationally exchanging the PI region of catalytically-“dead” Spy Cas9 (Spy dCas9) with those of the selected orthologs (Supplementary Figure S2A-B.^17,18^ Assembled variants, including Spy-mac dCas9, were separately co-transformed into *E coli* cells, along with guide RNA derived from *S. pyogenes* and an 8-mer PAM library of uniform base representation in the PAM-SCANR genetic circuit, established by others.^19^ The circuit usefully up-regulates a green fluorescent protein (GFP) reporter in proportion to PAM-binding strength. Therefore, we collected the GFP-positive cell populations by flow cytometry and Sanger sequenced them around the site of the PAM to determine position-wise base preferences in a corresponding variant’s PAM recognition. Spy-mac dCas9, more so than Spy-mut dCas9, generated a trace profile that was most consistent with guanine-independent PAM recognition, along with a dominant specificity for adenine dinucleotides (Figure 2A; Supplementary Figure S2C).

**Figure 2:**
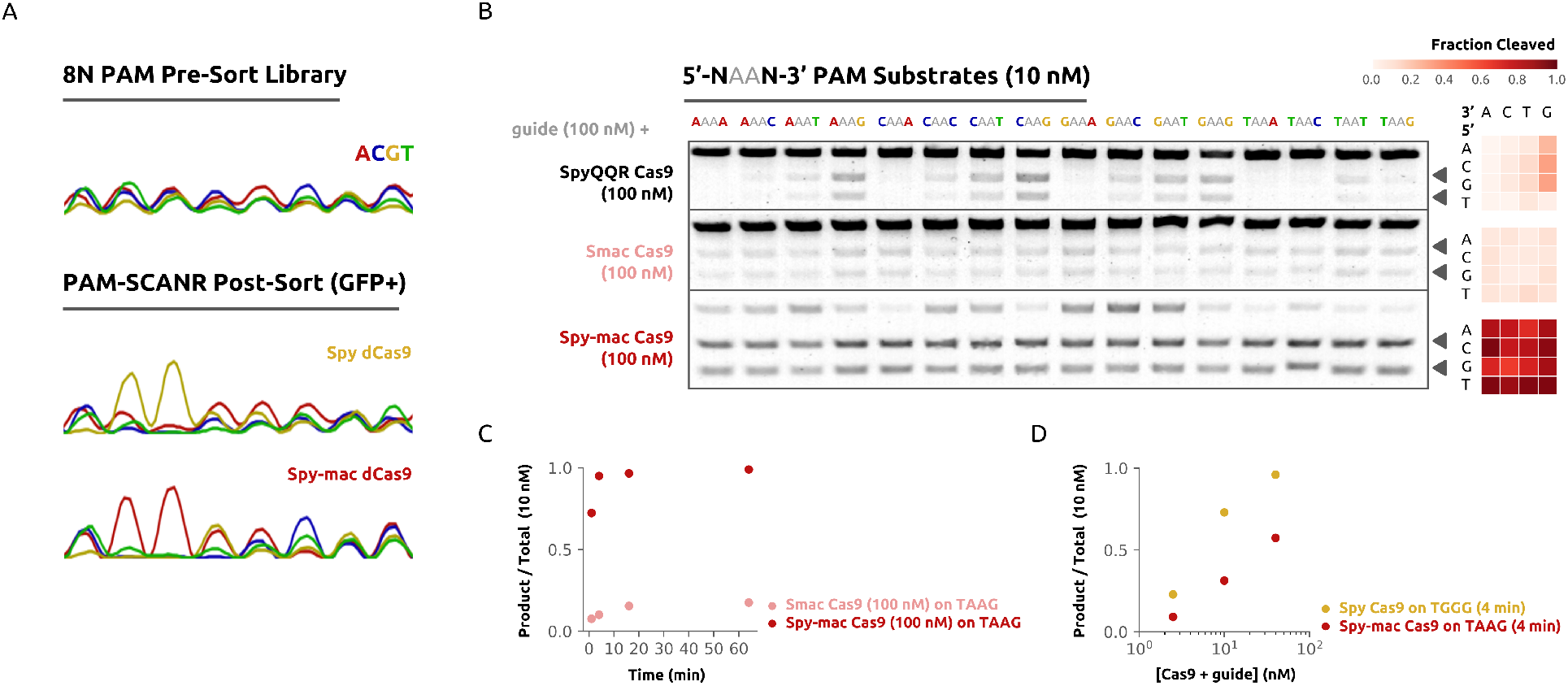
Validation of Smac Cas9 recognition for adenine dinucleotide PAM sequences. **(A)** Chromatograms representing the PAM-SCANR based enrichment of variant-recognizing PAM sequences from a 5’-NNNNNNNN-3’ library. **(B)** SYBR-stained agarose gels showing *in vitro* digestion of 10 nM 5’-NAAN-3’ substrates upon 16 minutes of incubation with 100 nM of purified ribonucleoprotein enzyme assemblies. Arrows distinguish banding of the cleaved products from uncleaved substrate (top band). Matrix plots summarize cleaved fraction calculations, which were carried out in a custom script for processing gel images. **(C)** Timecourse measurements of target DNA substrate cleavage for Smac Cas9 and Spy-mac Cas9. **(D)** DNA substrate cleavage plotted as a function of 0.25:1, 1:1, and 4:1 molar ratios of ribonucleoprotein to target for wild-type Spy Cas9 and hybrid Spy-mac Cas9.

Next, we purified nuclease-active enzymes to continue probing the DNA target recognition potential and uniqueness of Spy-mac Cas9 (Supplementary Figure S4A).^20,21^ We individually incubated the ribonucleoprotein complex enzymes (composed of Cas9 + crRNA + tracrRNA) with double-stranded target substrates of all 5’/3’-neighboring base combinations at an adenine dinucleotide PAM (5’-NAAN-3’; Figure 2B; Supplementary Table T1). Brief 16-minute digestion indicated both wild-type Smac Cas9 and the hybrid Spy-mac Cas9 cleaved adjacently to 5’-NAAN-3’ motifs more broadly and evenly than the previously reported QQR variant. Spy-mac Cas9 distinguished itself further with rapid DNA-cutting rates that resemble the fast digest kinetics of Spy Cas9 (Figure 2C-D).^22^ We ran reactions that used varying crRNA spacer lengths and tracrRNA sequence, as the latter differs slightly between the *S. macacae* and *S. pyogenes* genomes (Supplementary Figure S4B-E). Neither of these two parameters compensated for the slower cleavage rate of Smac Cas9, but we did notice marginal improvement in the activity of the wild-type form with its native tracrRNA, which comports with the interface of the guide-Cas9 interaction being mostly outside of the PI domain.

To verify that an adenine dsDNA dinucleotide is sufficient for Cas9 PAM recognition and target cleavage, we assembled target sequences that switch the next four downstream bases to the same nucleotide (e.g. 5’-TAAGXXXX-3’, for X all fixed to A, C, G, or T; Supplementary Figure S3F). We confirmed Spy-mac Cas9 remains active across this target set, albeit with some variation in cutting rate. Additionally, we observed moderate yield of cleaved products on examples of 5’-NBBAA-3’, 5’-NABAB-3’, 5’-NBABA-3’ PAM sequences (where B is the IUPAC symbol for C, G, or T; Supplementary Figure S4G), revealing an even broader tolerance for increments to the dinucleotide position or adenine adjacency. We anticipate future measurements of guide-loading, target-dissociation and R-loop expansion/contraction will provide more insights on the serendipitous catalytic benefit over Smac Cas9 from grafting its PI domain onto a truncated Spy Cas9.

Encouraged by the nucleolytic performance of Spy-mac Cas9, we investigated its capacity for gene modification in human cells (Supplementary Figure S5A). First, we transfected a human embryonic kidney (HEK293T) cell line with plasmids that encode Smac Cas9 or Spy-mac Cas9, and co-expressed single-guide RNA molecules that target the VEGFA gene locus at sites representing a breadth of 5’-NAAN-3’ PAM diversity (Supplementary Table T2). Consistent with *in vitro* observations, we found Spy-mac Cas9 was more efficient than Smac Cas9 at mediating enzymatically-detected (T7 EndonucleaseI) genomic insertion/deletion (indel) mutations. Spymac Cas9 also proved capable of generating indels with variable efficiency on instances of any directly 5’- or 3-neighboring base for 5’-NAAG-3’ or 5’-CAAN-3’ PAM sequences (Supplementary Figure S5B). To address sites with low modification rates, we introduced two mutations (R221K and N394K) into Spy-mac Cas9 that can raise gene knock-out percentages and had been previously identified by deep mutational scans of Spy Cas9.^23^ We refer to this variant as an “increased” editing Spy-mac Cas9 (iSpy-mac Cas9) due to its similarly elevated modification rates on most targets.

We then benchmarked the gene editing performance of the nucleases derived from *Streptococcus macacae* Cas9 against orthologs of Cas12a by making use of their common AT-rich PAM specificity.^24,25^ We included Cas12a orthologs known for efficient gene editing from *Acidaminococcus sp. BV3L6* (AsCas12) and *Lachnospiraceae bacterium ND2006* (LbCas12).^26^ Our selection of target sites permits overlapping PAM recognition between these Cas9 and Cas12a nucleases by guiding the Cas12a variants with the reverse complemented spacer sequences of those guiding Cas9 variants (Figure 3A). The Cas9 and Cas12a thereby targeted opposite strands, yet were constrained to recognize the same PAM site and preserve important features for guide RNA effectiveness (e.g. distribution of purines/pyrimidines, directionality of target-matching in relation to the PAM, and GC-content; Supplementary Table T2).^27,28^ We believe this is the first report of Cas12a and Cas9 activity being compared so explicitly on an endogenous genomic locus. For each site we examined, iSpy-mac Cas9 consistently generated a larger indel percentage than either AsCas12a or LbCas12a – never exhibiting less activity than the lower-editing of the two Cas12 proteins – if not generating the largest overall percentage (Figure 3B).

**Figure 3:**
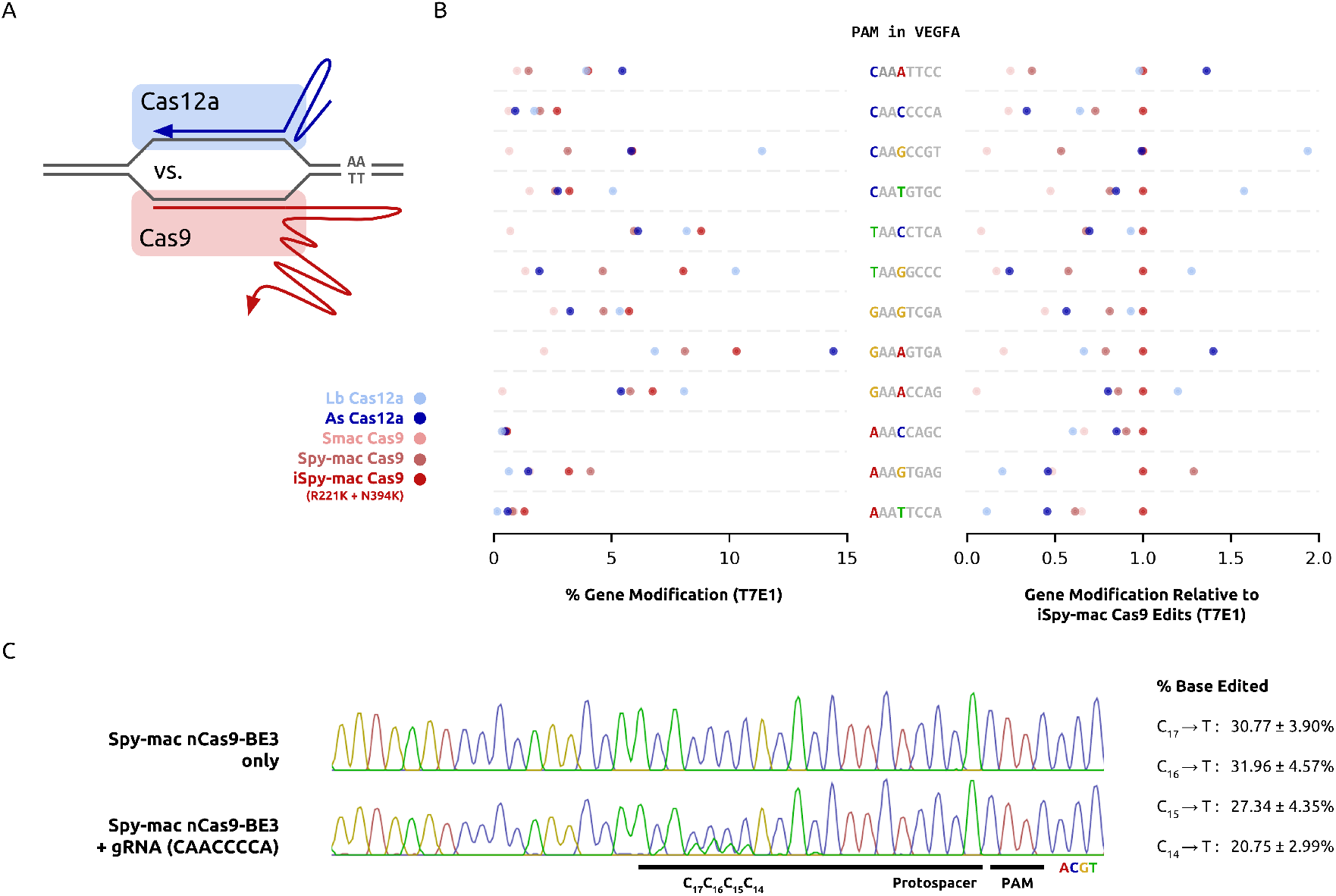
Confirmation of Spy-mac Cas9 as a unique and efficient genome engineering platform. **(A)** Schematic diagram for matching Cas9 and Cas12a guides in a manner that enforces their recognition of the same PAM sequence and therefore facilities their comparison (a “Cas12a vs Cas9 Comparator”). **(B)** Dot plots of absolute and relative gene modification efficiency in HEK293T cells by Cas9 and Cas12a variants targeting common PAM sequences located in the VEGFA gene. Values were quantified in a T7EI-based assay and are consistent with biological duplicates that were run in parallel. **(C)** Genomic base editing demonstration for the targeted conversion of cytosines to thymines with Spy-mac nCas9-BE3. Analysis on the efficiency was carried out in our custom Sanger sequencing trace file processing script called BEEP.

Lastly, we selected a window of four nucleotides in the VEGFA locus in a sequence context such that any other reported CRISPR endonuclease capable of gene modificiation would not allow their base editing with a cytidine deaminase-fused enzyme.^29^ Note, a Spy-mac Cas9 base editor has a distinct targeting range to implementations that use Cas12a since current base editing methods directly modify the non-target strand and in order to recognize the same PAM site, the two enzyme types must target in opposite orientations;^30,31^ Hence, Cas9 base editing architectures utilize their ability to nick on the guide-pairing target side of the R-loop structure (ribonucleoprotein bound and matched to DNA) to transfer a base edit in a manner that templates from the modified non-target strand.^32^ Accordingly, we co-transfected HEK293T cells with a nickase form of Spy-mac Cas9 derived from the previously reported BE3 architecture for cytosine base editing (Spy-mac nCas9-BE3) and the gRNA plasmid targeting a PAM downstream of the selected nucleotides.^5^ We measured robust levels of base editing in harvested cells, which exhibited 20% to 30% cytosine to thymine conversion at these positions (Figure 3C). Despite previous reports indicating base editing rates are generally lower than gene modification rates for the same target, we instead noticed a significant gain compared to the indel formation when we used double-strand breaking enzymes for this PAM site.^33^ Such discrepancy is likely explained by scaling to more sites for larger gene modification experiments, and possibly from differing codon usage outside of the PI domain. Recent work shows that higher editing rates can be achieved by optimizing such codon selection, nuclear-localization sequences/linkers, protein solubility, delivery methods, and sortable labeling of transfected cells.^34–37^

In summary, we have identified a homolog of Spy Cas9 in *Streptococcus macacae* with native 5’-NAAN-3’ PAM specificity. By leveraging the substantial background in the development and characterization of Spy Cas9, we engineered variants of Smac Cas9 that maintain its minimal adenine dinucleotide PAM specificity and achieve suitable activity for mediating edits on chromosomes in human cells.^38^ This finding sets the path for engineering enzymes like Spy-mac Cas9 with other desirable properties, control points, effectors, and activities.^33,39^–41 Spy-mac Cas9 can now open wide access to AT-content PAM sequences in the ever-growing list of genome engineering applications with Type-IIA CRISPR-Cas systems.

## Methods

### Selection of *Streptococcus* Cas9 Orthologs of Interest

All Cas9 orthologs from the *Streptococcus* genus were downloaded from the online UniProt database https://www.uniprot.org/. These were the downselected by pair-wise alignment to Spy Cas9 using a Blosum62 cost matrix in Genewiz software, discarding orthologs with less than 70% agreement with the Spy Cas9 sequence. The remaining 115 orthologs were used to generate a sequence logo (Weblogo http://weblogo.threeplusone.com/create.cgi), and were manually selected for divergence at positions aligned to residues critical for the PAM interaction of Spy Cas9. The SPAMALOT pipeline was implemented as we previously reported.^16^ Briefly, a set of scripts based around the Bowtie alignment tool (http://bowtie-bio.sourceforge.net) map the spacer sequences from CRISPR cassettes to putative targets in phage genomes. The SPAMALOT software can be downloaded at https://github.com/mitmedialab/SPAMALOT.

### PAM-SCANR Bacterial Fluorescence Assay

Sequences encoding the PAM-interaction domains of selected Cas9 orthologs were synthesized as gBlock fragments by Integrated DNA Technologies (IDT) and inserted via a New England Biolabs (NEB) Gibson Assembly reaction into the C-terminus of a low-copy plasmid containing Spy dCas9 (Beisel Lab, NCSU). The hybrid protein constructs were transformed into electrocompetent *E. coli* cells with additional PAM-SCANR components as previously established.^19^ Overnight cultures were analyzed and sorted on a Becton Dickinson (BD) FACSAria machine. Sorted GFP-positive cells were grown to sufficient density, and plasmids from the pre-sorted and sorted populations were then isolated. The region flanking the nucleotide library was PCR amplified and submitted for Sanger sequencing (Genewiz). The choromatograms from received trace files were inspected for post-sorted sequence enrichments relative to the pre-sorted library.

### Purification of and DNA cleavage with Selected Nucleases

The gBlock (IDT) encoding the PAM-interaction domain of *S. macacae* was inserted into a bacterial protein expression/purification vector containing wild-type *S. pyogenes* Cas9 fused to the His6-MBP-tobacco etch virus (TEV) protease cleavage site at the N-terminus(pMJ915 was a gift from Jennifer Doudna, Addgene plasmid #69090). The resulting hybrid Spy-mac Cas9 protein expression construct was sequence-verified by a next-generation complete plasmid sequencing service (CCBI DNA Core Facility at Massachusetts General Hospital). The hybrid-protein construct was then transformed into BL21 Rosetta 2^TM^(DE3) (MilliporeSigma), and a single colony was picked for protein expression, inoculated in 1 L 2xYT media, and grown at 37 Celsius to a cell density of OD600 0.6. The temperature was then lowered to 18 Celsius and His-MBP-TEV-SpyMac Cas9 expression was induced by supplementing with 0.2 mM IPTG for an additional 18 hours of growth before harvest.

Cells were then lysed with BugBuster^TM^Protein Extraction Reagent, supplemented with 1 mg/ml lysozyme solution (MilliporeSigma), 125 Units/gram cell paste of Benzonase^TM^Nuclease (MilliporeSigma), and complete, EDTA-free protease inhibitors (Roche Diagnostics Corporation). The lysate was clarified by centrifugation, including a final spin with a pre-chilled Steriflip^TM^0.45 micron filter (MilliporeSigma). The clarified lysate was incubated with Ni-NTA resin (Qiagen) at 4 Celsius for 1 hour and subsequently applied to an Econo-Pac^TM^chromatography column (Bio-Rad Laboratories). The protein-bound resin was washed extensively with wash buffer (20 mM Tris pH 8.0, 800 mM KCl, 20 mM imidazole, 10% glycerol, 1 mM TCEP) and His-tagged Spy-mac protein was eluted in wash buffer (20 mM HEPES, pH 8.0, 500 mM KCl, 250 mM imidazole, 10% glycerol). ProTEV^TM^Plus protease (Promega, Madison) was added to the pooled fractions and dialyzed overnight into storage buffer (20 mM HEPES, pH 7.5, 500 mM KCl, 20% glycerol) at 4 Celsius using Slide-A-Lyzer^TM^dialysis cassettes with a molecular weight cut-off of 20 KDa (ThermoFisher Scientific). The sample was then incubated again with Ni-NTA resin for 1 hour at 4 Celsius with gentle rotation and applied to a chromatography column to remove the cleaved His tag. The protein was eluted with wash buffer (20 mM Tris pH 8.0, 800 mM KCl, 20 mM imidazole, 10% glycerol, 1 mM TCEP) and fractions containing cleaved protein were verified once more by SDS-PAGE and Coomassie staining, then pooled, buffer exchanged into storage buffer, and concentrated. The concentrated aliquots were measured based on their light-absorption (Implen Nanophotometer) and flash-frozen at –80 Celsius for storage or used directly for *in vitro* cleavage assays.

The crRNA and tracrRNA guide components were procured in the form of HPLC-purified RNA oligos (IDT) and resuspended in 1X IDTE pH 7.5 solution (IDT). Duplex crRNA-tracrRNA guides were annealed at 1 uM concentration in duplex buffer (IDT) by a protocol of rapid melting followed by gradual cooling. Target substrates were PCR amplified from assemblies of the PAMSCANR plasmid with a fixed PAM sequence. *In vitro* digestion reactions with 10 nM target and typically a 10-fold excess of enzyme components were prepared on ice and then incubated in a thermal cyclcer at 37 Celsius. Reactions were halted after at least 1 minute of incubation by subsequent heat denaturation at 65 Celsius for 5 minutes and run on a 2% TAE-agarose gel stained with DNA-intercalating SYBR dye (Invitrogen). Gel images were recorded from blue-light exposure and analyzed in a Python script adapted from https://github.com/jharman25/gelquant/. Cleavage fraction measurements were quantified by the relative intensity of substrate and product bands as follows: 
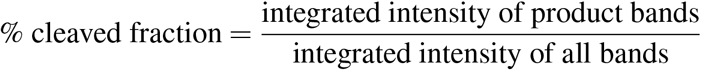

### Gene Modification Analysis and Software

The gBlock (IDT) encoding the PAM-interaction domain of *S. macacae* was swapped into the Spy Cas9 mammalian expression plasmid OG5209 (Oxford Genetics). Plasmids for Cas12a protein plus Cas9 and Cas12a guide construction were gifts from Keith Joung (Addgene plasmid 78741, 78742, 78743, 78744). HEK293T cells were maintained in DMEM supplemented with 100 units/ml penicillin, 100 mg/ml streptomycin, and 10% fetal bovine serum (FBS). sgRNA plasmid (62.5 ng) and nuclease plasmid (187.5 ng) were transfected into cells as duplicates (5 × 10^4^/well in a 96-well plate) with Lipofectamine 3000 (Invitrogen) in Opti-MEM (Gibco). After 5 days post-transfection, genomic DNA was extracted using QuickExtract Solution (Epicentre), and genomic loci were amplified by PCR utilizing the KAPA HiFi HotStart ReadyMix (Kapa Biosystems). For indel analysis, the T7EI reaction was conducted according to the manufacturer’s instructions and equal volumes of products were analyzed on a 2% agarose gel stained with SYBR Safe (Thermo Fisher Scientific). Gel image files were analyzed in a Python script adapted from https://github.com/jharman25/gelquant/. Boundaries of cleaved and uncleaved bands of interest were hard-coded for each duplicate set of Cas variants with a common target, and the areas under the corresponding peaks were measured and calculated as the fraction cleaved of the total product. Percent gene modification was calculated as follows: 
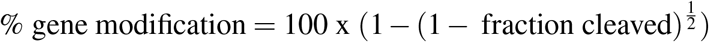

### Base Editing Analysis and Software

The gBlock (IDT) encoding the PAM-interaction domain of *S. macacae* was swapped into a mammalian expression plasmid for cytosine to thymine base editing, which came as a gift of David Liu (Addgene plasmid 73021). HEK293T (ATCC^®^*[circlecopyrt]* CRL-3216^TM^) cells (MilliporeSigma, Burlington, MA) were maintained in DMEM supplemented with 100 units/ml penicillin, 100 mg/ml streptomycin, and 10% fetal bovine serum (FBS). sgRNA (500 ng) and BE3 plasmids (500 ng) were transfected into cells as duplicates (2 × 10^5^/well in a 24-well plate) with Lipofectamine 3000 (Invitrogen) in Opti-MEM (Gibco). After 5 days post-transfection, genomic DNA was extracted using QuickExtract Solution (Epicentre), and the VEGFA genomic locus was amplified by PCR utilizing the KAPA HiFi HotStart ReadyMix (Kapa Biosystems). Amplicons were purified and submitted for Sanger sequencing (Genewiz). For base conversion analysis, an automated Python script termed BEEP, employing the pandas data manipulation library and BioPython package, was utilized to align base-calls of an input ab1 file to first determine the absolute position of the target within the file, and subsequently measure the peak values for each base at the indicated position in the spacer. Finally, editing percentages of specified base conversions were calculated and normalized to that of an unedited control. Conversion efficiencies are reported as the average of two independent duplicate reactions ± standard deviation. The BEEP software can be downloaded at https://github.com/mitmedialab/BEEP.^16^

## Author Contributions

N.J. and P.C. conceived identification strategies for PAM novelty, designed and implemented workflows for PAM discovery, and conducted data analysis for PAM validation. N.J. identified Smac Cas9 and related orthologs as proteins of interest. L.N. assembled ortholog constructs for PAM characterizations, optimized protein purification protocols, and isolated nucleases for enzymology. N.J. and P.C. formulated and carried out experiments to evaluate genome editing in mammalian cell culture. J. M. J. supervised the study. All authors contributed to writing and editing the manuscript.

## Acknowledgments

We thank Professor Neil Gershenfeld and Dr. Shuguang Zhang for shared lab equipment. We thank Professor Ed Boyden for access to tissue culture.

## Correspondence

Correspondence and requests for materials should be addressed to N.J.

---

**Figure S1.**
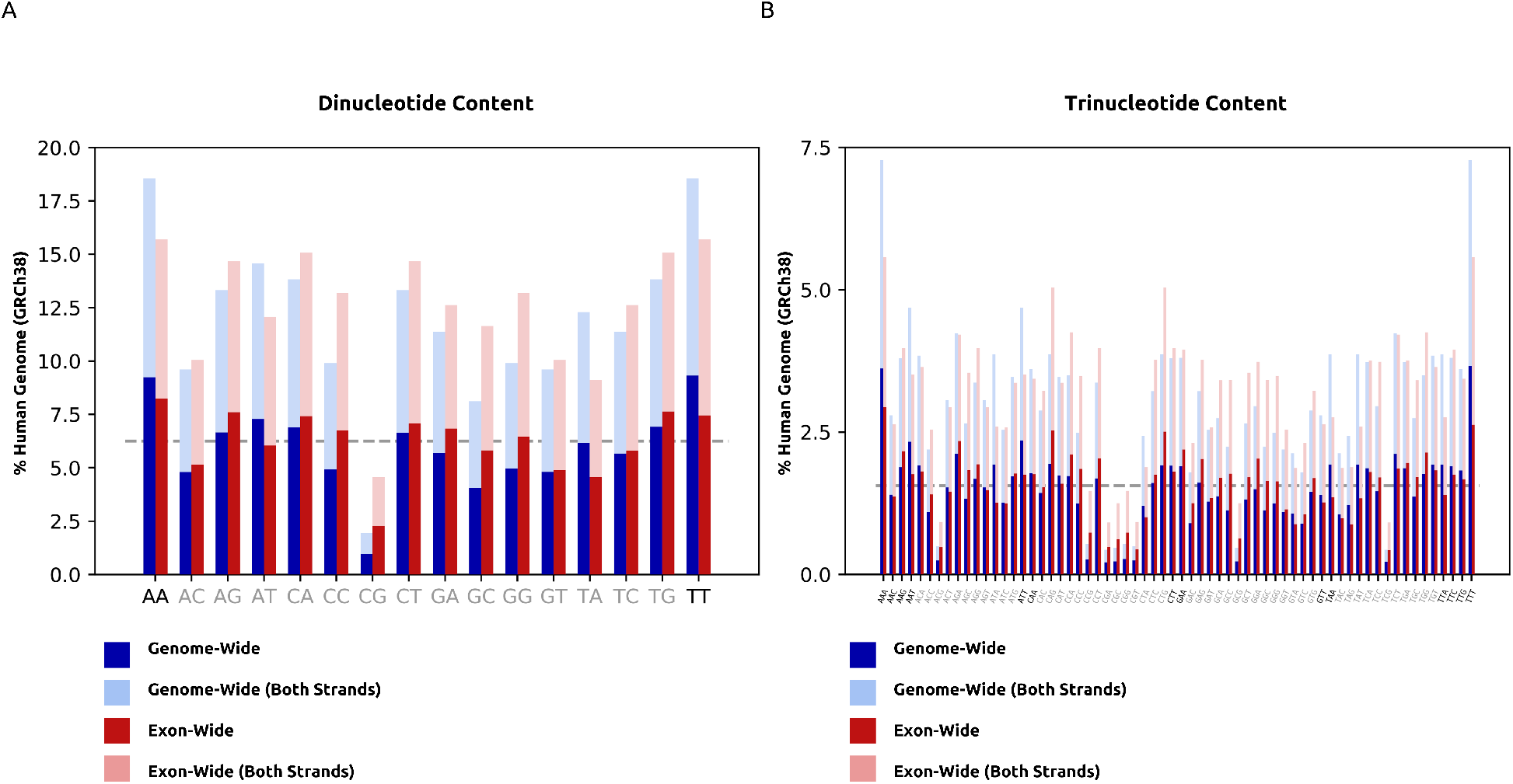
**(A)** Dinucleotide and **(B)** Trinucleotide occurrences in the human reference genome GRCh38. Tallies were carried out using the compseq EMBOSS command line software tool. Dashed gray lines mark what the expected percentages would be for a uniform representation of all sequences of length 2 or 3.

**Figure S2.**
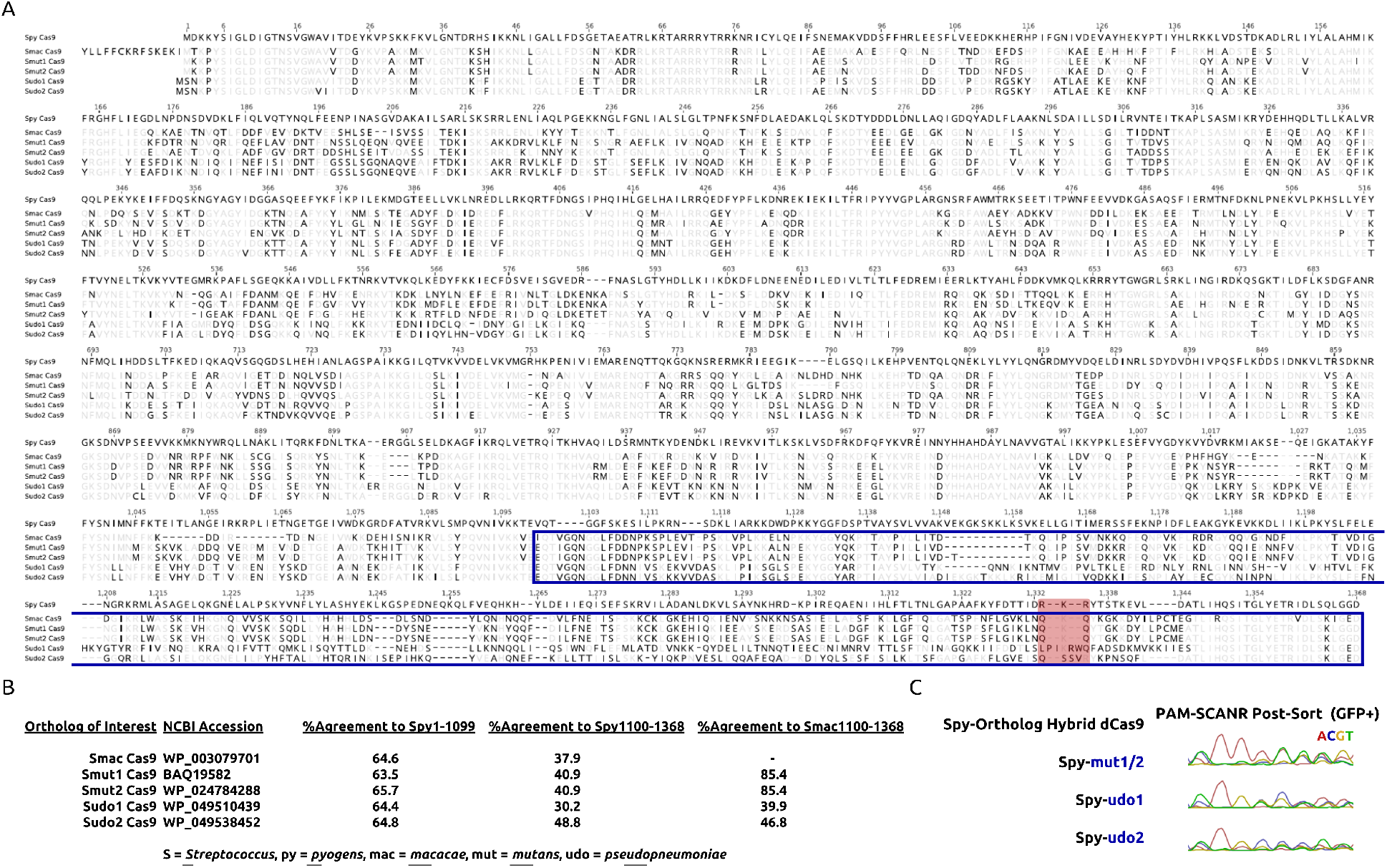
**(A)** Sequence alignment (Genewiz software) for selected orthologs of interest that substitute at least one critical PAM-contacting arginine residue within the region highlighted in red. A blue box marks the C-terminal component grafted onto truncated Spy Cas9 to form dCas9 hybrids. **(B)** Table listing the homology shared within and outside of the box to those regions in the corresponding Spy Cas9 and Smac Cas9 reference sequences. **(C)** Chromatograms representing the PAM-SCANR based enrichment of variant-recognizing PAM sequences from a 5’-NNNNNNNN-3’ library.

**Figure S3.**
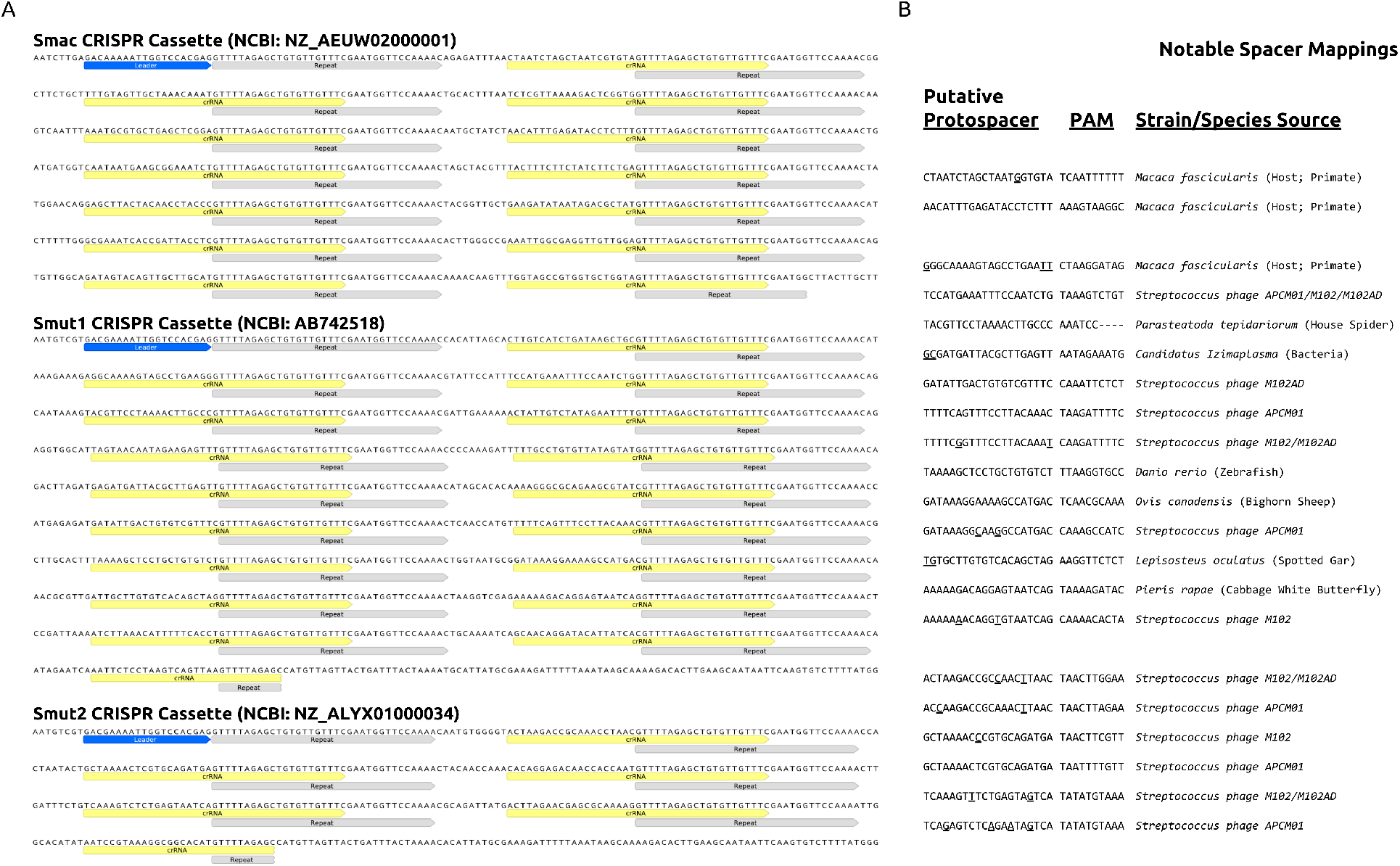
**(A)** Annotated CRISPR cassettes obtained from the genomes corresponding to orthologs that substitute both PAM-contacting arginine residues to glutamine. **(B)** Mappings of CRISPR cassette spacers to their putative target source for listed crRNA, identified via an online BLAST and/or SPAMALOT. SPAMALOT uncovered most cases of mismatch-tolerated mappings to *Streptococcus phage*. Underlined bases indicate mismatches that are tolerated for the mapping. Additional line spacing separates analysis for each CRISPR cassette.

**Figure S4.**
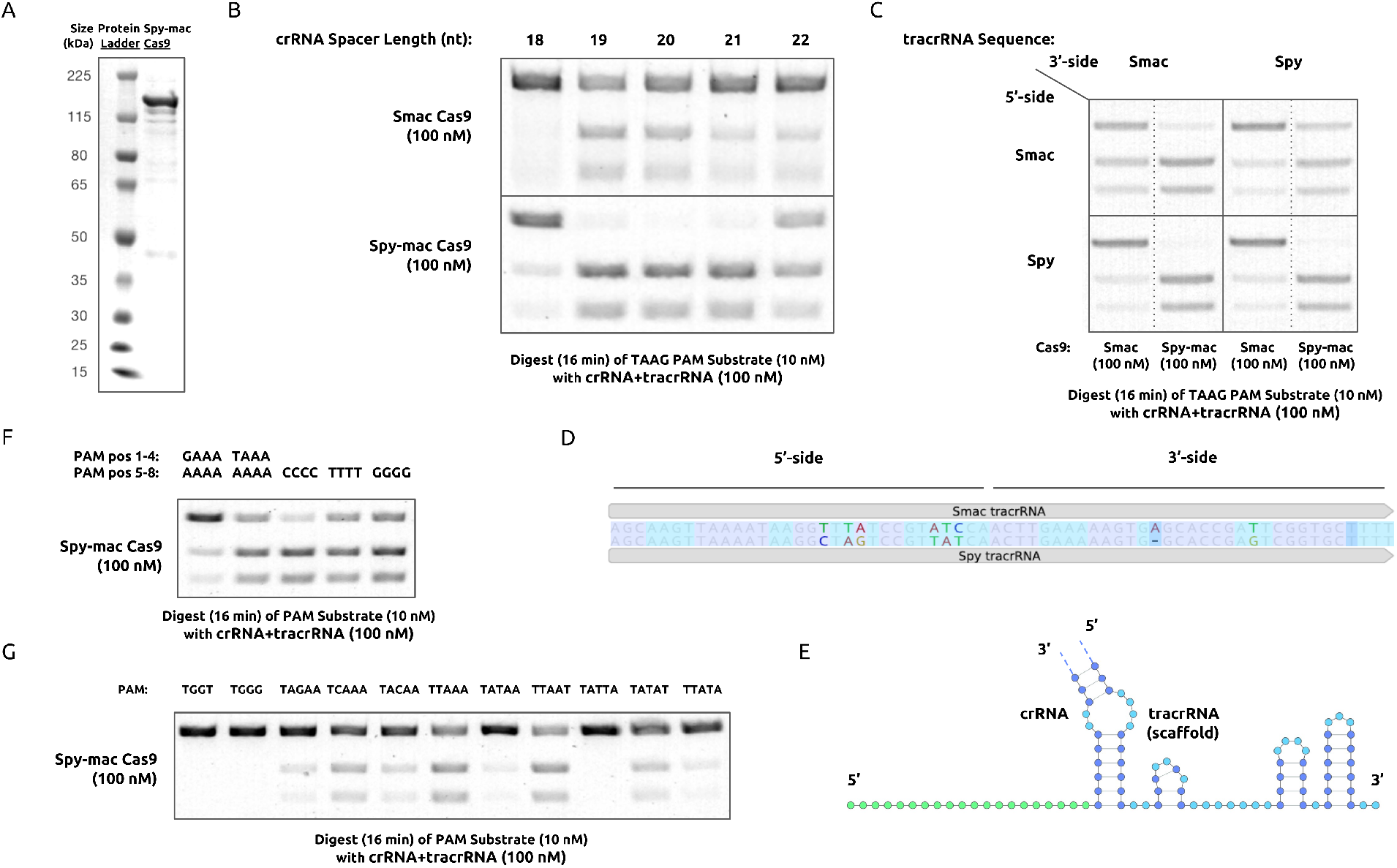
**(A)** SDS-PAGE gel image of Spy-mac Cas9 after purification by affinity chromatography. SYBR-stained agarose gels running *in vitro* digestion reactions are shown that assay dependencies on **(B)** crRNA spacer length and **(C)** tracrRNA sequence origin. **(D)** Sequence alignment (Genewiz software) of tracrRNA from *S. pyogenes* and *S. mutans* highlighted in a color code that reflects the base-pairing in their **(E)** duplex gRNA secondary structure. SYBR-stained agarose gels running *in vitro* digestion reactions are shown that assay dependencies on **(F)** positions 5–8 in the PAM sequence and **(G)** increments to the distribution of adenine content in positions 1–5 in the PAM sequence.

**Figure S5.**
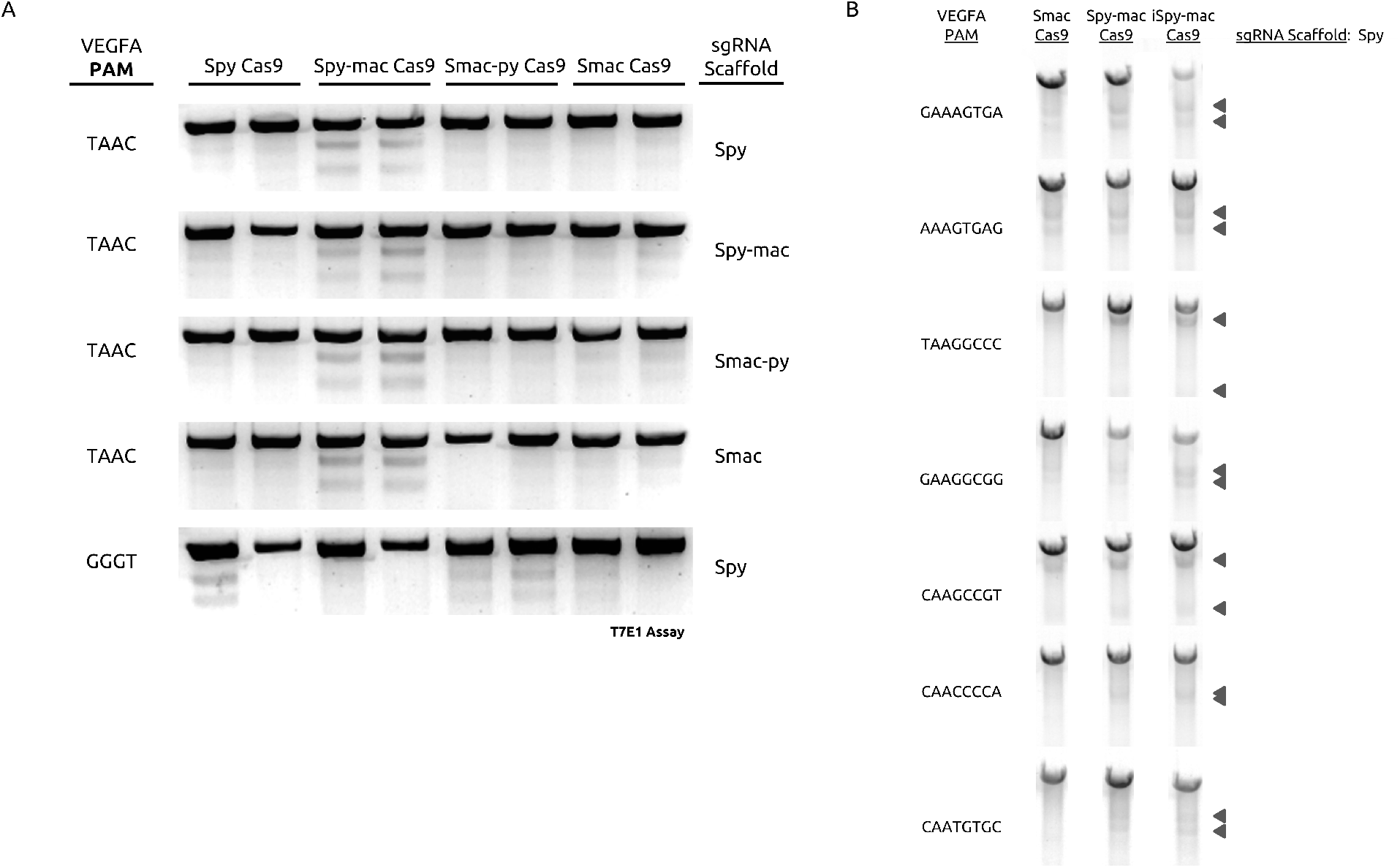
Detection of genomic modification in SYBR-stained agarose gels running T7EI digests upon targeting **(A)** a single PAM site with combinations of wild-type plus hybrid variants of Cas9 and guide scaffold (tracrRNA sequence) from *S. pyogenes* and *S. macacae*, and **(B)** a diversity of PAM sequences with the wild-type and engineered variants that include the Smac Cas9 PI domain. Arrows point to the banding from products digested by T7EI, which is used to estimate gene modification efficiencies.

**Table T1.**
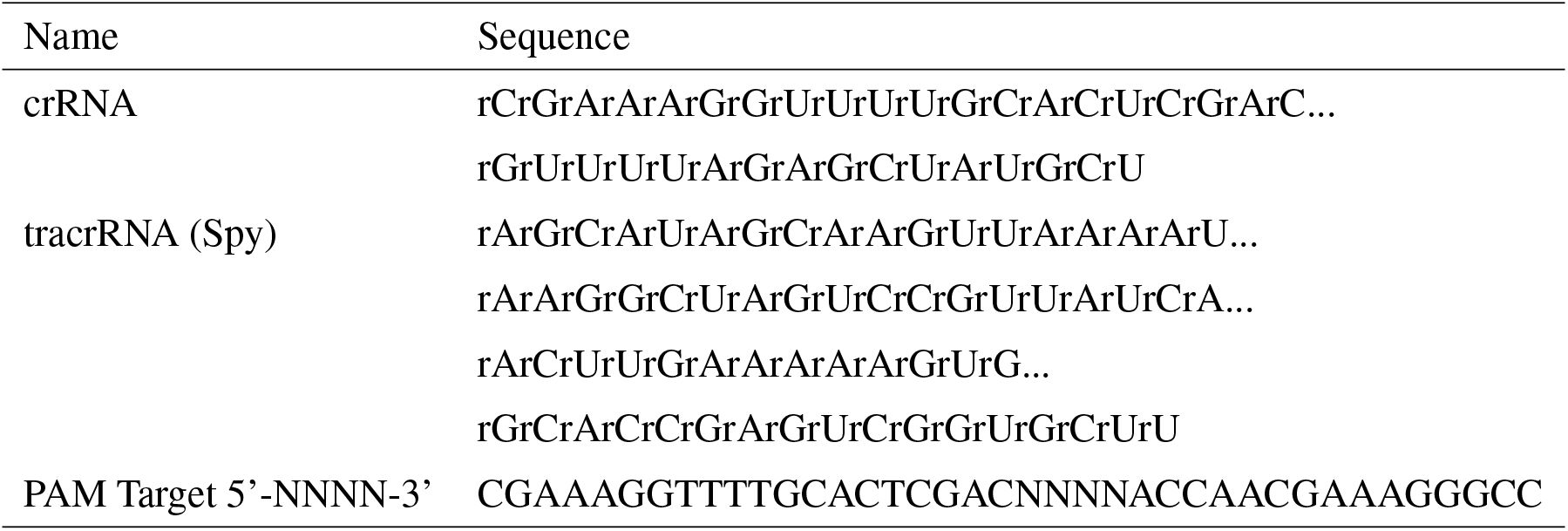
Sequence information for *in vitro* digest reactions

**Table T2.**
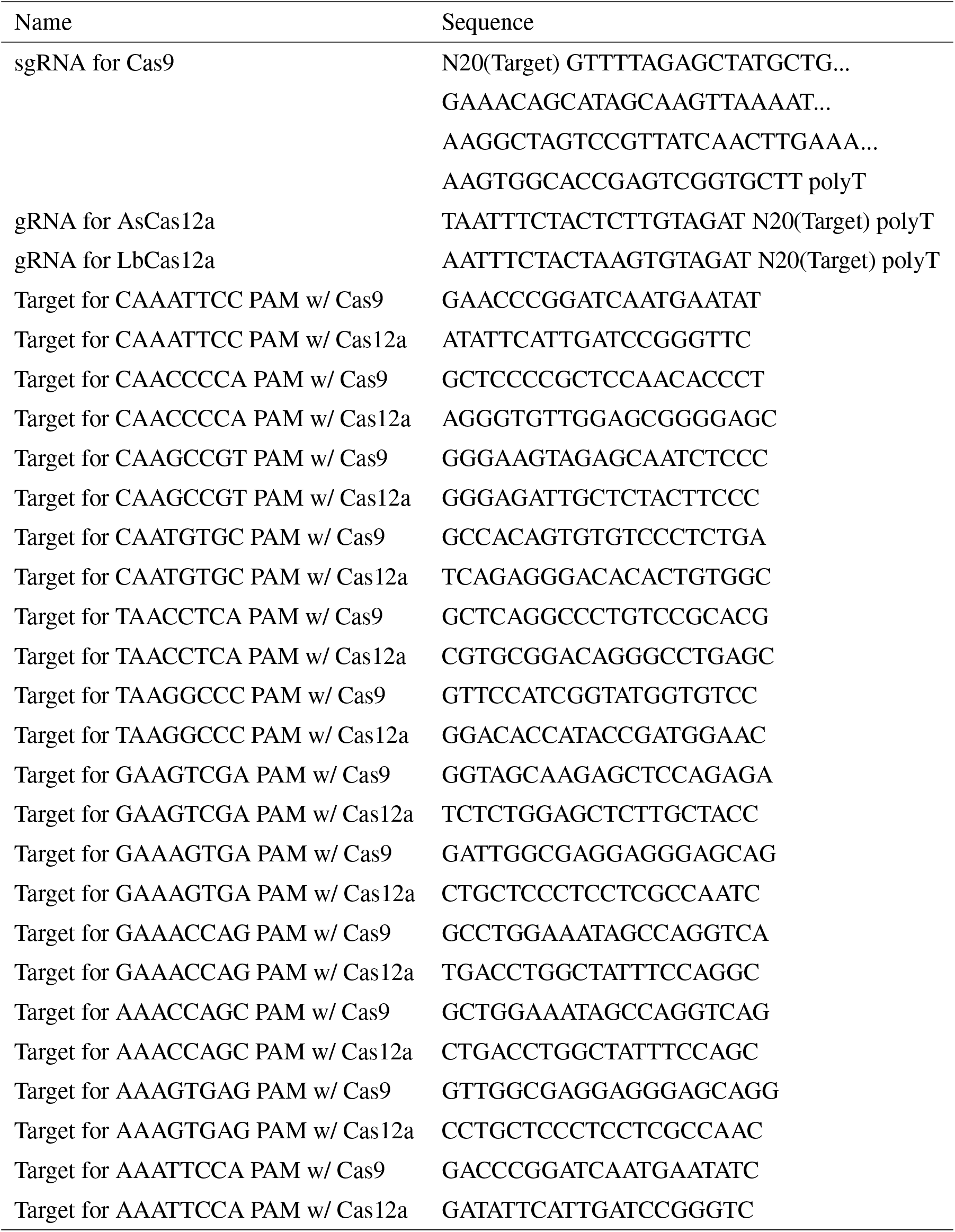
Sequence information for genome editing in human cells

